# Life Identification Numbers: A bacterial strain nomenclature approach

**DOI:** 10.1101/2024.03.11.584534

**Authors:** Federica Palma, Melanie Hennart, Keith A. Jolley, Chiara Crestani, Kelly L. Wyres, Sebastien Bridel, Corin A. Yeats, Bryan Brancotte, Brice Raffestin, Sophia David, Margaret M. C. Lam, Radosław Izdebski, Virginie Passet, Carla Rodrigues, Martin Rethoret-Pasty, Audrey Combary, Solene Cottis, Martin C. J. Maiden, David M. Aanensen, Kathryn E. Holt, Alexis Criscuolo, Sylvain Brisse

## Abstract

Unified strain taxonomies are needed for the epidemiological surveillance of bacterial pathogens and international communication in microbiological research. Core genome multilocus sequence typing (cgMLST) holds great promise for standardized high-resolution strain genotyping. However, this approach faces challenges including classification instability and disconnection of new nomenclature from widely adopted classical MLST identifiers. This essay discusses the cgMLST-based Life Identification Number (LIN) method, recently proposed as a stable multilevel strain taxonomy system applicable to most bacterial pathogens. We describe how LIN codes are implemented and used in practice for precise strain definitions and epidemiological tracking.

**Glossary:** 

**Multilocus sequence typing (MLST):** A genotyping method applied mostly to microbial strains to study population structure and epidemiology, based on comparing the nucleotide sequences of a small number (typically seven) of housekeeping protein-coding genes. In MLST, allele numbers are assigned to each sequence variant (allele) of a given gene. The MLST genotype of a bacterial strain is defined by the combination of the allele numbers observed at the genes that are included in the genotyping scheme. A sequence type (ST) is assigned to each unique combination of alleles, called an MLST profile. MLST was invented in 1998 and became a de-facto standard taxonomy of bacterial strains, albeit at low resolution.

**Core genome MLST:** An extension of MLST that analyzes sequence variation across hundreds to thousands of conserved (core) genes, shared by all strains of a species, providing higher resolution typing for genomic epidemiology and evolutionary studies. cgMLST schemes typically comprise 2000 to 4000 genes, depending on the genome size and genetic variation (in terms of presence/absence of genes) within bacterial species. A core genome sequence type (cgST) can be assigned to unique cgMLST profiles, i.e., a unique combination of cgMLST allelic numbers.

**Whole Genome Sequencing (WGS):** A method that determines the complete DNA sequence of an organism’s genome in a single process, providing comprehensive information for comparative genetic analyses based on cgMLST or other analytic methods.

**Single Nucleotide Polymorphisms (SNPs):** Variations at a single base position in the DNA sequence among individuals isolates, strains or species, used as genetic markers for studying for example, evolutionary relationships or strain identity.

**Average nucleotide identity (ANI):** A measure of genomic similarity between two organisms, calculated as the average percentage of identical nucleotides in orthologous genomic regions; commonly used to assess species-level relatedness in prokaryotes.

**Taxonomy:** Here, we apply the word taxonomy to bacterial strains as a system of classifying, naming and identifying strains based on shared genetic characteristics as defined by *e.g.,* cgMLST.

## Bacterial strain taxonomies: why and how?

Taxonomies of bacterial strains responsible for infectious diseases are essential resources to ensure effective communication in population biology, epidemiological surveillance, and public health response to outbreaks. As illustrated by the SARS-CoV-2 variant nomenclature, simple nicknames (*e.g.,* Alpha, Delta, Omicron can greatly improve communication among multiple actors, including the public, in face of public health threats (1,2). Strain taxonomies are therefore needed to precisely recognize and define variants with properties of special medical interest, such as antimicrobial resistance, high virulence or vaccine escape.

Taxonomic systems are based on three pillars: classification, nomenclature and identification. Currently, there are neither classification nor nomenclature standards to define sublineages, variants, types or clones (hereafter, collectively called “strains”) within bacterial species (3). Linnean taxonomy encompasses classification levels from Phylum down to subspecies, but the latter is seldom used for bacterial species as it is not a practical solution to describe strains. Ad-hoc phenotypic (*e.g.,* serotypes) and genotypic (*e.g.,* sequence types) methods have long been used to differentiate strains from particular species but have shown limitations in terms of universal applicability, reproducibility of classification, or level of resolution. However, the advent of universally applicable whole genome sequencing (WGS) has advanced the potential to refine and generalize strain taxonomy by providing the maximal discrimination power needed for epidemiological surveillance, while being broadly applicable as a harmonized approach across pathogen Phyla (4–6). Yet, few attempts have been made to devise genomic taxonomies of strains and evaluate their general applicability. With WGS now being implemented worldwide in all sectors of microbiology (*e.g.*, medical, veterinary, food, environmental), a precise and universal procedure for describing bacterial strains becomes a key need to translate WGS data into relevant information that would support epidemiological surveillance, outbreak investigations, cross-niche or between-host transmission tracking, and public health actions that need international and cross-sectoral coordination.

Several bacterial strain taxonomies have emerged in recent years. One of them, applied to bacterial pathogens with low amounts of genetic diversity (*i.e.,* evolutionary recent pathogens), relies on recognizing notable branches in phylogenetic trees, defined by specific diagnostic single nucleotide polymorphisms (7,8). A similar approach, the Pango nomenclature, was successfully applied to SARS-CoV-2 variants (2). Unfortunately, these phylogenetic approaches face challenges raised by the need to update phylogenetic trees and define novel lineages. Another classification approach, PopPunk, relies on pairwise comparisons of unaligned genomes (based on *k*-mers) to create groups within bacterial populations, and is scalable to large and diverse datasets (9). However, among the broad range of methods developed for bacterial strain typing and group naming (10,11), multi-locus sequence typing (MLST), based on the analysis of a few (typically seven) conserved loci, was established over the last two decades as the method of choice for most bacterial species (12–14). Indeed, major strengths of MLST are its standardization, as it relies on well-defined fixed sets of genetic markers, and its ease of interpretation and portability. Classical MLST schemes form the basis of widely adopted “sequence type” (ST) taxonomies (15) in most bacterial species, which are maintained, expanded and made available to the international community through the platforms BIGSdb (16) and EnteroBase (17).

The logical extension of MLST at genome scale, known as core genome MLST (cgMLST), uses thousands of conserved gene loci, leading to the definition of core genome sequence types, or cgST (4,18). However, because of their much higher resolution, any cgST will match only with a tiny fraction of bacterial isolates from a given bacterial species, making the cgST a less useful nomenclatural element than the classical ST for tracing genetic relationships through broad space and time scales. To define phylogenetic associations among similar cgSTs, which together might represent meaningful groups of particular medical or epidemiological interest (a common way of conceptualizing the informal notion of ‘strain’), cgMLST allele profiles can be grouped at chosen levels of dissimilarity, resulting in multilevel classifications. A single-linkage clustering was initially used to create these higher-level groups from cgMLST data (e.g., (19)), but by design this approach suffers from a lack of stability, as preexisting groups can merge when intermediate genotypes are sampled. To address this issue, profiles can be assigned to the most closely related preexisting group. This approach, called Hierarchical clustering (HierCC), represents the first multilevel bacterial strain taxonomy system based on cgMLST (20). HierCC is implemented in the platform EnteroBase, where taxonomies for strains of *Salmonella*, *Escherichia coli* and other important bacterial pathogens are maintained (17,21).

Amongst the various efforts to align the taxonomy of all life forms with genomic data, a novel multilevel classification system was proposed by Vinatzer and colleagues, using multi-position numerical codes attributed to each individual genome (22,23). These codes, called Life Identification Numbers (LINs), were designed to encompass all domains of life in a single, unified taxonomy, based on the Average Nucleotide Identity (ANI) metric (24). A database of these ANI-based LIN codes, GenomeRxiv (initially called LINbase; https://genomerxiv.cs.vt.edu), was set up to enable global development and use of this taxonomic approach (25).

Given that the pairwise ANI estimates between genomes can be imprecise for nearly identical strains, particularly when draft genomes are significantly fragmented, some of us proposed combining LIN codes with the cgMLST approach in order to design novel taxonomies of bacterial strains within species (26). We found the use of pairwise dissimilarities between cgMLST profiles, rather than ANI estimates between genomes, provides greater reproducibility in appraising small-scale genome relationships. Hence, cgMLST-based LIN codes (hereafter, LIN codes for short) combine the strengths of both approaches.

Here, we first present this LIN code approach and its recent improvements, including (i) its implementation in the widely used genotyping platforms BIGSdb and Pathogenwatch (16,27) and (ii) a LIN code nicknaming procedure to facilitate the designation of familiar intra-species groups of key importance in either biological research or epidemiological surveillance. Next, we illustrate how the LIN codes can be used to address questions in population biology and genomic epidemiology, using the case of the *Klebsiella pneumoniae* Species Complex (KpSC), a phenotypically and genetically diverse ubiquitous pathogenic group (28). This essay underlines the benefits of LIN codes for stable definition and labelling of intra-species groups from epidemiologically important phylogenetic lineages down to outbreak strains.

## cgMLST-based LIN coding, and missing data handling

LIN codes are series of numeric codes that reflect genomic similarity between organisms. cgMLST-based LIN code systems consist of multiple (*e.g.*, 10) predefined positions (or bins), each corresponding to a range of pairwise cgMLST profile similarity values, together representing a partition of the complete range [0%-100%]. From left to right, the positions of the code correspond to decreasing allele mismatch dissimilarity, *i.e*., increasing similarity. The leftmost bins are thus used to classify deep phylogenetic divisions, whereas the rightmost bins will distinguish recently evolved variants. Analogous to the classification levels of Linnean taxonomy (*e.g.*, Phylum – Class - Order – Family – Genus – species), the LIN codes thus capture from left to right, the membership of a genome to taxonomic groups of increasing relatedness.

The process of LIN code assignment from cgMLST data, first proposed in Hennart *et al*. (26), is summarized in **Figure 1** (and formalized in **Section 1** of the Supplementary appendix). A LIN code is created for each distinct cgST. The system is initialized by creating, for a selected initial cgST, a LIN code with the integer value 0 at every bin. The initial cgST can be chosen randomly or using a reference strain of the species or group under consideration (*e.g.*, the first strain that was sequenced, or the taxonomic type strain). The next steps are the same for each subsequent individual cgST. Each incoming cgST is matched against all already LIN-encoded cgSTs, in order to identify its most similar one (hereafter, the reference cgST), based on the fraction of allele mismatches across cgMLST loci. For creating the novel LIN code, the pivot bin is defined as the bin in which the observed allele similarity falls, and the novel LIN code is then created in three steps (**Figure 1**): (i) copying the LIN code prefix of the reference cgST, *i.e.* from the leftmost bin up to the pivot bin (excluded); (ii) incrementing by 1 the maximum integer value observed in the pivot bin among the cgST(s) sharing the same prefix used at step (i); (iii) attributing the integer value 0 at the bins downstream of the pivot, corresponding to initialization of the novel subdivision created at the pivot bin level.

**Figure 1.**
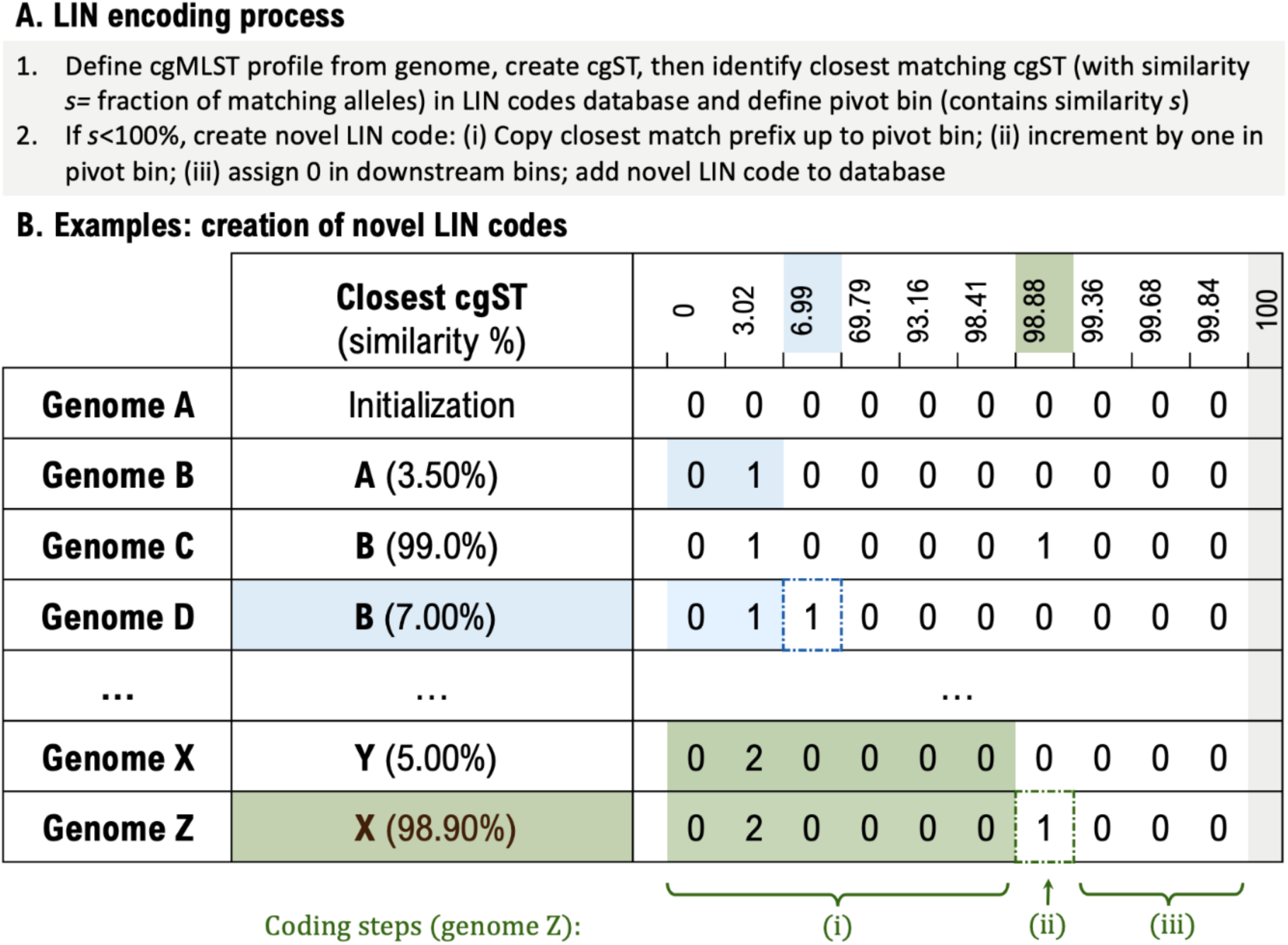
Overview of the process of cgMLST-based LIN code assignment. **A. LIN encoding process**. The process starts with assigning cgMLST profiles to genome sequences and next with classifying profiles into unique core genome sequence types (cgST). One genome, and its associated cgST, is selected to initiate the process, with 0 assigned to each bin. For each incoming genome, the closest already encoded cgST is then identified (based on fraction of matching alleles) and its similarity recorded to define the pivot bin (if the similarity is 100%, the LIN code is simply assigned to the query cgST, but no novel LIN code is created). When the similarity is <100%, a novel LIN code is created following steps (i), (ii) and (iii) (see details in main text and in the Supplementary Appendix). **B**. **Examples of novel LIN code creation**. The similarity threshold values given in the header line correspond to those defined for the KpSC LIN code scheme (see correspondence with minimal allelic mismatches and other information in the Supplementary appendix). Note that there is no bin corresponding to complete similarity (gray column on the right), as in this case the LIN codes are identical, *i.e*., there is no need to create a novel LIN code. Technically, each bin has a left border threshold (inclusive) that corresponds to a maximum number of pairwise allele differences between profiles, and is delimited on the right by the next threshold (exclusive, as the threshold value corresponds to the left threshold of the downstream bin – or, for the last bin, corresponds to LIN code identity). The first row (Genome A) corresponds to the unique initialization step (full-0 code for the initial cgST). Note that similarity values defining the bins each correspond to fixed numbers of shared alleles among cgMLST profiles, divided by the length of the cgMLST scheme.

The cgSTs are different from the classical ST in an important way: due to the often-fragmented nature of genome sequence assemblies and because many core genes are dispensable in bacteria, the cgST must accommodate missing data. If two cgSTs (new and reference) have a 100% similarity (*i.e.*, no allele mismatch among the loci called in both profiles), the LIN code of the reference is simply assigned to the new cgST. This can happen when the new cgST differs from the reference only by its missing data pattern, such pairs (or groups) being called coincident cgSTs (see Supplementary appendix, **Section 2**). Consequently, a single LIN code can correspond to multiple coincident cgSTs.

An important implication of the encoding process is that LIN codes are created definitively, as are the assignments for LIN codes to individual cgSTs. Hence, LIN codes are stable by design, and the incorporation of novel genomes will never affect pre-existing LIN codes and their assignments. This provides trust in LIN code referencing and stability in comparisons across time.

### The internal structure of LIN codes and the notion of prefix

A LIN code prefix can be defined as any bin subset that starts from the leftmost position. An important particularity of LIN codes is that the numerical identifiers at a given bin position (except the leftmost one) can only be interpreted in the context of the LIN code prefix preceding it: the same integer value at a given bin position corresponds to group membership only if the upstream prefixes are identical. In other words, the integer values at a given bin position are subdivisions of their respective upstream prefixes, and their numbering starts from zero independently for each prefix. This minimizes the maximal value used in each bin making them easy to read. This property can be regarded as a systematization of the analogous possibility in the Linnean nomenclature, where the same species epithet can be used for distinct genera (*e.g*., *Klebsiella pneumoniae* and *Streptococcus pneumoniae*). The initialization at zero for prefix subdivisions contrasts with other taxonomic systems, such as the hierarchical clustering approach used in the genomic epidemiology platform EnteroBase (17,21), in which a group identifier is created independently at each level (see Supplementary appendix, **Section 3**, for details).

The notion of shared LIN code prefix is also important because it conveys a sense of genetic similarity among genomes: the longer the common prefix of two LIN codes is, the more similar the two corresponding genomes (in the strict sense, cgMLST profiles). For a given cgST profile, its LIN code thus expresses how similar it is to every other genome in the LIN code taxonomy. Very different profiles will show identity at few or no prefix positions of their LIN codes, whereas nearly identical genomes will yield LIN codes identical at most or all prefix positions (see *e.g.,* **Figure 1**, genomes Z *versus* X: the shared prefix 0_2_0_0_0_0 implies a minimum similarity of 98.88% (inclusive) and a maximum similarity of 99.36% (exclusive). We note that our definition of LIN code prefix is similar to the LINgroup concept proposed by Vinatzer and colleagues (23).

### Nicknaming LIN code prefixes provides continuity with previous nomenclatures

Whereas LIN code prefixes themselves can serve as machine-readable ‘diagnostic’ markers of groups of interest, they are not very easy to remember or pronounce by humans. It was therefore proposed to nickname relevant LIN code prefixes with simple denominations (23). A prefix nicknaming system was also implemented within the BIGSdb platform for cgMLST-based LIN codes. It is thereby possible to nickname every distinct prefix in any chosen way. For example, one option is to increment an integer identifier (analogous to the numbering of STs in the MLST framework) for each novel prefix of a given length; but alternative labelling could be applied, such as Greek letters, astronomical objects, or any other series of words that may be universally understandable and easy to remember. This nicknaming process would be particularly useful for long prefixes, or prefixes of phenotypic or taxonomic relevance that subdivide the population at particularly informative levels.

For bacterial species with established nomenclatures, nicknames can be assigned to LIN code prefixes based on prior denominations such as MLST or serotyping, to retain interpretation and recognition as much as possible. To enable backward compatibility of LIN codes with well-established ST identifiers, a majority identifier inheritance rule was developed (26). For example, in the KpSC LIN code system, prefixes are nicknamed using ST identifiers as a source (the process is formally defined in the Supplementary appendix, **Section 8**). For convenience, groups of KpSC genomes with the same prefix of length 3 or 4 were designated as sublineages (SL) or clonal groups (CG), respectively (see **Figure 2**). The prefixes corresponding to these two levels were nicknamed because they correspond to deep subdivisions of the KpSC population structure, and their partitions (*i.e*., single prefixes) are highly concordant with well-known MLST-based STs (Supplementary appendix, **Section 11**).

**Figure 2.**
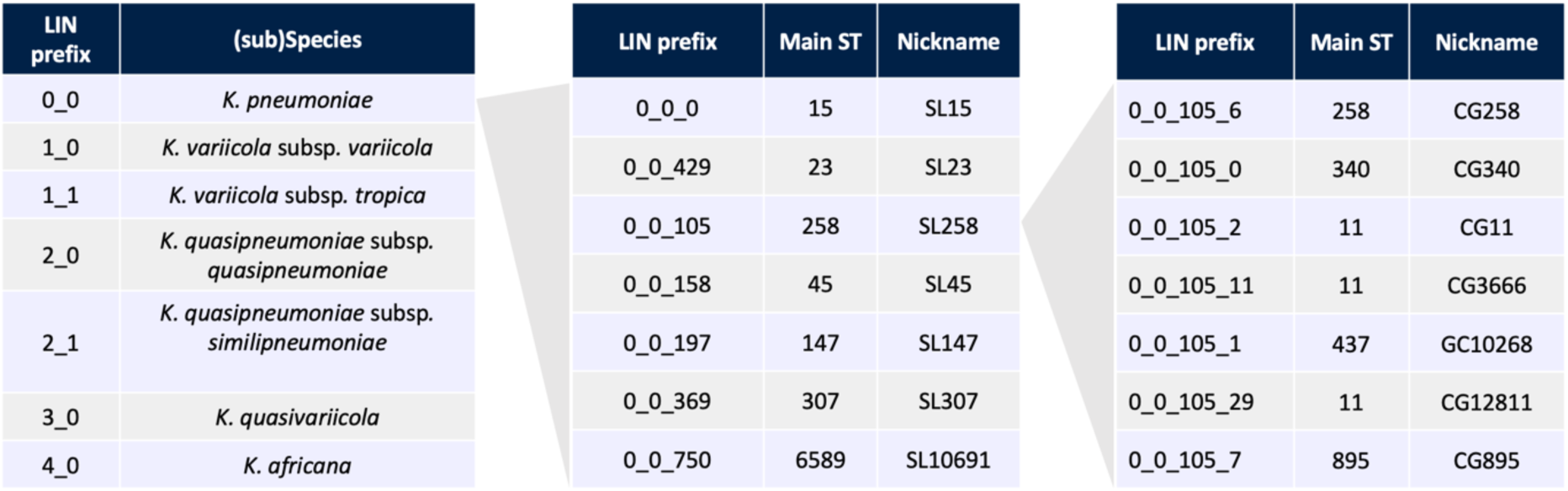
Nicknaming of LIN code prefixes enables inheritance of previous nomenclatures. Nicknames of some KpSC LIN code prefixes of lengths 2 to 4 bins, inherited from Linnaean taxonomy (2-bin prefix, left panel) or 7-gene MLST (prefixes of lengths 3 and 4 bins, central and right panels, respectively). Groups corresponding to level 3 are called sublineages (SL) and groups of level 4 are called clonal groups (CG).

Although KpSC ST identifiers and SL/CG nicknames are generally identical, some ST numbers are shared by phylogenetically distinct genomes that can result from recombination events leading to the same combination of the seven alleles (see Supplementary appendix, **Section 11**). Therefore, prioritization of LIN code nicknames over classical STs, is recommended in future.

### How to design LIN codes: population structure and outbreak datasets as a guide

An initial requirement for any cgMLST-based LIN code taxonomy implementation is a cgMLST scheme that has been designed and thoroughly validated to ensure the stability of the taxonomy from its inception. Subsequent removal of loci (e.g., because they are too infrequently called), or changing their template (e.g., choosing an upstream start codon) would result in potential inconsistencies with previously defined LIN codes. Therefore, it is advisable to define LIN code taxonomies following careful evaluation of the cgMLST schemes from which they are derived, particularly if these are intended for broad usage.

The scope of applicability of the cgMLST scheme is also an important consideration. MLST or cgMLST schemes are typically used for a single species, and less frequently for an entire genus (*e.g*., more than 90% ANI). In rare cases, an intermediate category called species complex is covered; these correspond to groups of closely related species that are sometimes misidentified in routine microbiology diagnostic processes. The applicability of the cgMLST scheme (and related LIN code taxonomy) should ideally be broadened, but increasing the phylogenetic breadth will be at the expense of the core genome size, reducing the discriminatory power of the scheme.

While any number of bins (up to the number of loci in the cgMLST scheme) can be chosen to create a LIN code system, it is recommended to guide their definition by analyzing the population structure of the species, in order to propose phylogenetically informative bin thresholds. Several methods have been designed to find optimal ranges of dissimilarities that optimize the reliability of the subsequent classifications (20,26,29).

Deep levels might also correspond to previously recognized subdivisions and might therefore be optimized to match these previous classifications. For example, in the KpSC scheme, the distinction of its phylogroups (taxonomic species or subspecies) was used as a guide to define the two deepest thresholds (26), as detailed in Supplementary appendix, **Section 7**.

For epidemiological levels, bin thresholds can be selected to reflect epidemiological surveillance practice. In a hypothetical example, four different alleles might be typically used to define clusters and trigger outbreak investigations; in this case, using a LIN code bin associated with a threshold of four allele differences would be congruent with this practice. Broader epidemiological thresholds might yield more false positives (sporadic isolates unrelated to a given common source outbreak) in detecting genetic clusters, but will on the other hand capture outbreaks during which more diversity has accumulated. It is therefore advised to use a set of epidemiological thresholds which will be useful in different situations, from the most stringent (*i.e.*, one single allelic difference) to more relaxed ones.

### Visualization tools for LIN codes as proxies of population diversity

Once LIN codes are implemented and genomes encoded, the repertoire of LIN codes can be used to derive summary views of the species diversity and its structure. The complete and up-to-date LIN code nomenclature (comprising alleles, profiles, cgSTs and LIN codes) can be extracted from BIGSdb using a single query; for example for the KpSC, the LIN code taxonomy is available at https://bigsdb.pasteur.fr/api/db/pubmlst_klebsiella_seqdef/schemes/18/profiles_csv.

LIN code diversity within a bacterial species can be summarized using circular packing plots, which illustrate the diversity of populations at each classification level (**Figure 3)**. These representations also convey a sense of relative frequencies of the variants and enable the identification of the most epidemiologically represented populations.

**Figure 3.**
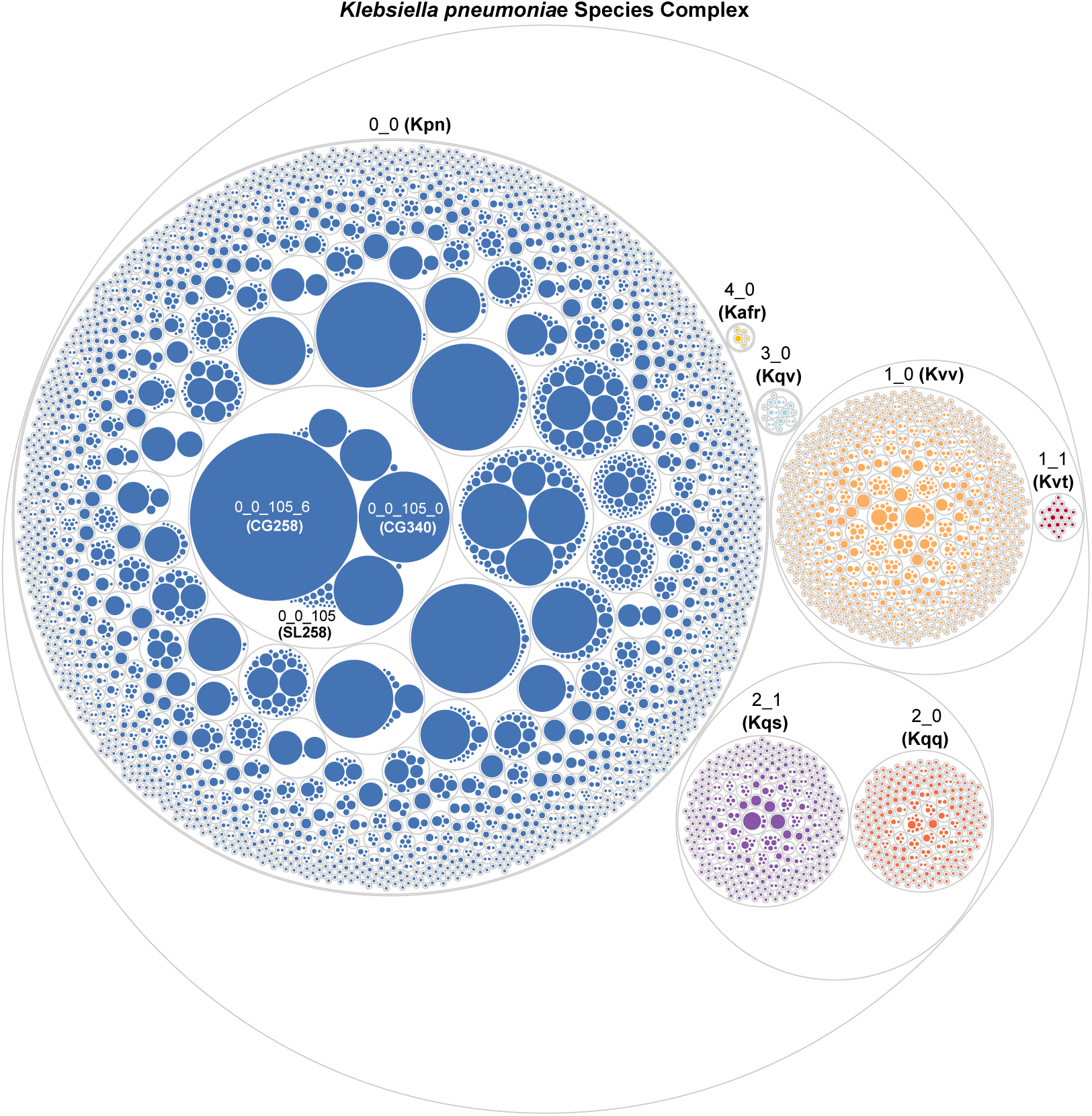
The hierarchical nature of LIN code positions applied to KpSC. The hierarchical structure of LIN codes is shown via a circular packing plot (data from the BIGSdb-Pasteur KpSC database). The circles correspond to LIN code prefixes of lengths 1 to 4 (an extra, all-encompassing circle corresponds to the entire KpSC); the size of each circle is related to the number of genomes it comprises. Numbering starts from 0 for subdividing each higher-level partition, characterized by a unique LIN code prefix. The first two bins in the LIN codes are used to identify Linnaean taxa. Whereas for three species there is a unique 2-bin prefix (*e.g.*, prefix 0_0 for *K. pneumoniae* [Kpn], 3_0 for *K. quasivariicola* [Kqv], 4_0 for *K. africana* [Kafr]), in the other cases two subspecies are distinguished (2_0 for *K. quasipneumoniae* subsp. *quasipneumoniae* [Kqq] and 2_1 for *K. quasipneumoniae* subsp. *similipneumoniae* [Kqs]; 1_0 for *K. variicola* subsp. *variicola* [Kvv] and 1_1 for *K. variicola* subsp. *tropica* [Kvt]). The hierarchical nature of LIN codes applies to subsequent levels such as those corresponding to sublineages (third bin, *e.g.* Kpn SL258 is identified with the LIN code prefix 0_0_105) and to clonal groups (fourth bin, *e.g*. the LIN code prefix 0_0_105_6 corresponds to Kpn CG258). Data was plotted using ggplot2 (R v4.3.2) and edited using Inkscape.

Second, the nested structure of a set of LIN codes can also be visualized by representing the associated prefix tree, which roughly approximates the phylogenetic relationships among isolates (26); see Supplementary appendix, **Section 6**. In such prefix trees, each internal node corresponds to a distinct LIN code prefix, where each node’s height corresponds to the associated bin threshold. This tree topology can be built without the initial genomes or cgMLST profiles, based solely on the relationships encoded in the nomenclature itself, and can serve as a computationally light proxy for the phylogenetic relationships among isolates.

### LIN codes in practice: source databases of taxonomies, and their use with external tools

A taxonomic system needs to be created and updated in a coordinated manner. The cgMLST LIN code strain taxonomy approach was first implemented in BIGSdb, since v1.34.0 (26). This open-source application is so far the only platform that has implemented cgMLST-based LIN codes, and is deployed at two main sites, PubMLST (at Oxford University) and BIGSdb-Pasteur (at Institut Pasteur, Paris). For the KpSC, the BIGSdb-Pasteur database serves as the source database for the definitions of alleles, cgMLST profiles, cgSTs, and LIN codes. For other pathogens, the PubMLST platform is the source database of LIN code taxonomies (see Future directions and conclusions).

A LIN code taxonomy is created with reference to a defined indexed scheme (*i.e.*, a scheme with a unique identifier for each profile, *e.g.,* cgST), with allele mismatch thresholds that define the LIN code bins. **Figure 1** and **Section 7** in the Supplementary appendix give details for the KpSC example of a LIN code taxonomy.

Compared to the stand-alone tools initially used to create cgMLST-based LIN code taxonomies (26), the implementation of LIN codes into the BIGSdb application has been accompanied by a number of important improvements, including (i) ensuring the reproducibility of LIN encoding by addressing the dependency of this approach to rounded genetic distance values (Supplementary appendix, **Section 4**); (ii) implementing input order rules for creating novel LIN codes (Supplementary appendix, **Section 5**); and (iii) implementing formal rules for handling missing data (as described above; Supplementary appendix, **Section 2**). These improvements were introduced to achieve the robustness needed for a reference taxonomy. Furthermore, functionality was developed in BIGSdb for searching database isolates based on LIN code or prefixes (Supplementary appendix, **Section 9**).

Currently, there is no stand-alone bioinformatics workflow to generate and handle local LIN code taxonomies. Although managing local LIN code taxonomies might be attractive for confidentiality reasons, this usage would be restricted to internal comparisons and would not fulfil the intended shared nomenclature objective of open, central LIN code taxonomies.

To make the LIN code taxonomy broadly accessible, its components (alleles, profiles, cgSTs, LIN codes and nicknames) can be extracted from BIGSdb source taxonomy databases using an application programming interface (30). They can then be used with external tools and analysis platforms, for example by extracting cgMLST alleles from local genome sequences and matching these with the source nomenclature data (**Figure 4**). As a first example of external use of LIN codes, we implemented LIN code matching functions within the Pathogenwatch platform, which supports KpSC genomic typing (27) and now matches genome sequences with an internal copy of the KpSC reference LIN code taxonomy (Supplementary appendix, **Section 10**).

**Figure 4.**
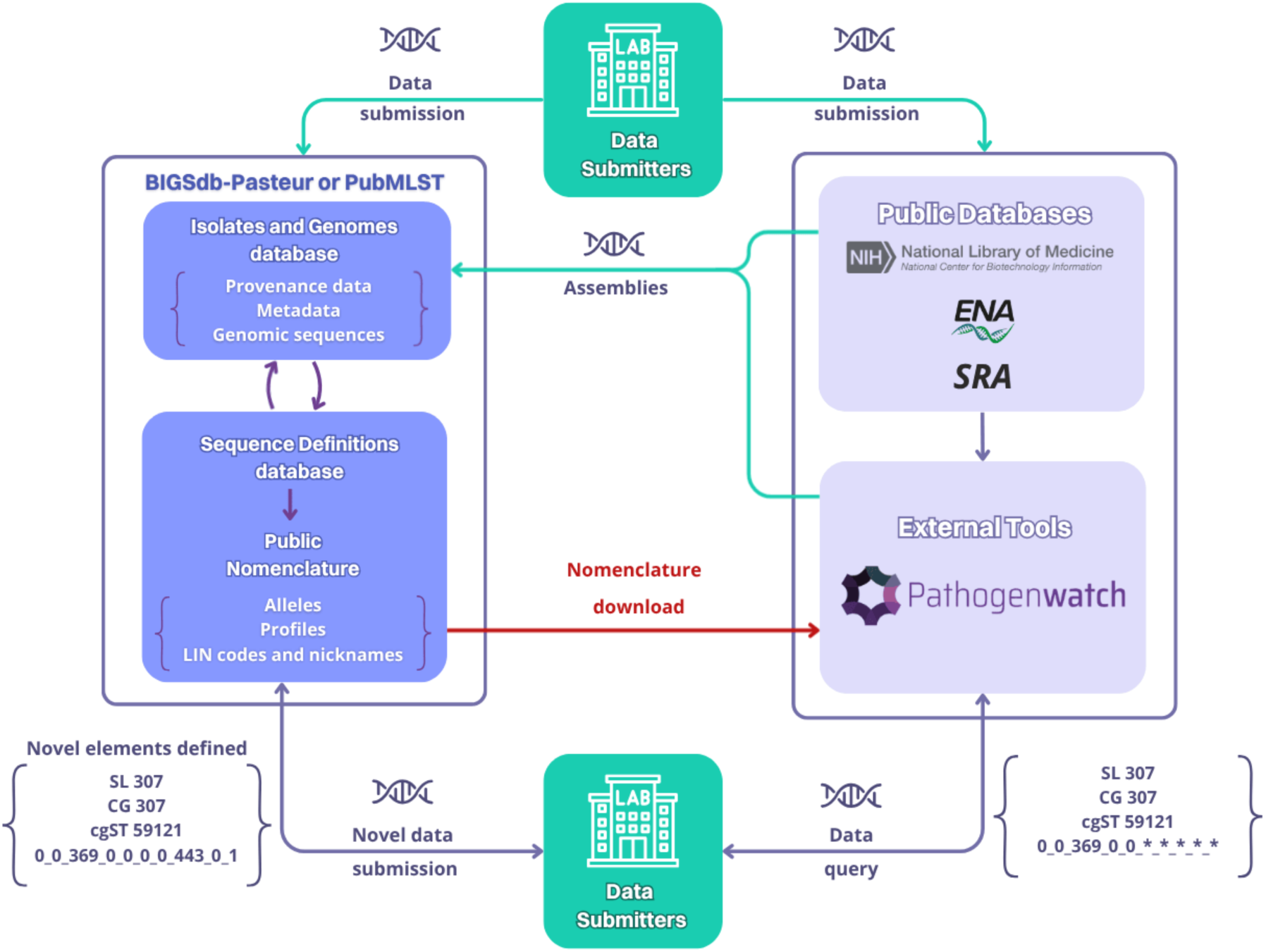
The LIN code taxonomy ecosystem. The source database of LIN code taxonomy (‘Sequence Definitions database’, lower left dark blue box) hosts the taxonomic elements (alleles, profiles, LIN codes). Curators create taxonomic elements from data sourced directly from data submitters, from NCBI/ENA, or from Pathogenwatch assemblies derived from SRA short-read data (green arrows). External tools or platforms such as Pathogenwatch can retrieve the LIN code taxonomy from BIGSdb using Application Programming Interfaces (API; red arrow), such that query genome sequences can be compared to the copy of the reference taxonomy in order to define their closest match (blue arrow, bottom right). The LIN code bins that can be defined are then reported (followed by asterisks for undefined ones), as well as sublineage and clonal group nickname information (if this can be extracted from the deduced LIN code). In this example, although the LIN code is incomplete, the genome can be inferred as belonging to clonal group 307 (defined as prefix 0_0_369_0). To obtain a complete LIN code, the genomic sequence (or its extracted taxonomic elements) must be submitted to the source database (blue arrow, left) so that novel taxonomic elements can be defined consistently.

When novel genome sequences are matched to the LIN code taxonomy, no identical cgMLST profile may exist at that time in the source LIN code taxonomy, implying that the cgST and complete LIN code cannot be determined. Still, the level of similarity between the query genome and the closest reference cgMLST profile enables inference of their common prefix; in other words, the LIN code of the query genome can be partially defined. If the query genome is closely related to one in the nomenclature database, its LIN code will be almost completely defined. Hence, the use of LIN codes in external databases or tools can have great functional relevance.

However, in the most general case, an incomplete match will be found. This implies that new nomenclatural elements (cgST profiles and LIN codes) have been discovered and could be defined for the benefit of the global community. This can only be done within the source database, otherwise the consistency of nomenclature will be lost. For any genome that has no complete LIN code, data submission to the source database is therefore encouraged. Furthermore, to be effective, external copies of the LIN code database need to be frequently (*e.g*., daily) synchronized with the primary database, given that the latter is updated continuously.

### LIN code applications in epidemiological surveillance and outbreak investigations

Bacterial species can harbor huge amounts of genetic diversity and are often structured genetically into recognizable sublineages. For example, in *K. pneumoniae*, sublineages including SL258, SL147, SL307, SL17 and SL23 have been recognized as globally distributed drivers of multidrug and/or hypervirulent infections. These sublineages have been the subject of detailed studies, that have led to defining their geographical spread and phylogenetic subgroups (31–35). However, so far, these sublineages have been defined using a mix of 7-locus MLST, cgMLST, and ANI; a clear and simple definition, and a harmonized nomenclature, have been lacking, making it difficult to recognize them in subsequent studies.

One prominent example of how LIN codes provide clear definitions of sublineages and disambiguate MLST definitions, is the case of hypervirulent sublineage ST23. Whole genome sequence analyses demonstrated the polyphyletic status of ST23 (Lam et al., 2023), which conflates isolates from two distant phylogenetic branches, which are appropriately separated into two LIN code sublineages (SL23: 0_0_429 and SL218: 0_0_115; Supplementary appendix, **Section 11**). Here we show that beyond the case of ST23, multiple distinct sublineages are conflated into single STs, but that they are also appropriately recognized by their distinctive LIN codes (Supplementary appendix, **Section 11; Table S1**).

We further illustrate how LIN codes can help track dissemination at fine genetic scales within sublineages, using the example of SL258, a major *Klebsiella pneumoniae* carbapenemase (KPC)-producing sublineage of *K. pneumoniae*. SL258 is defined by the LIN code prefix 0_0_105 and encompasses all isolates from 7-gene ST11, ST258, ST340, ST512 and some others (see **Figure 2**). Its phylogenetic structure is depicted in **Figure 5** (see Supplementary appendix, **Section 12** for methodological details) and shows that SL258 is divided into several subclades. These include CG258 (0_0_105_**6**), which contains all ST258 and ST512 isolates. LIN code bin 5 can further be used to distinguish major subclades within SL258, including those corresponding to ST340 (0_0_105_**0**_**11**), ST437 (0_0_105_**1**_**1**) and other subclades within ST11, some of which appearing to be associated with recombination events that include the capsule (K) locus (KL column in **Figure 5**).

**Figure 5.**
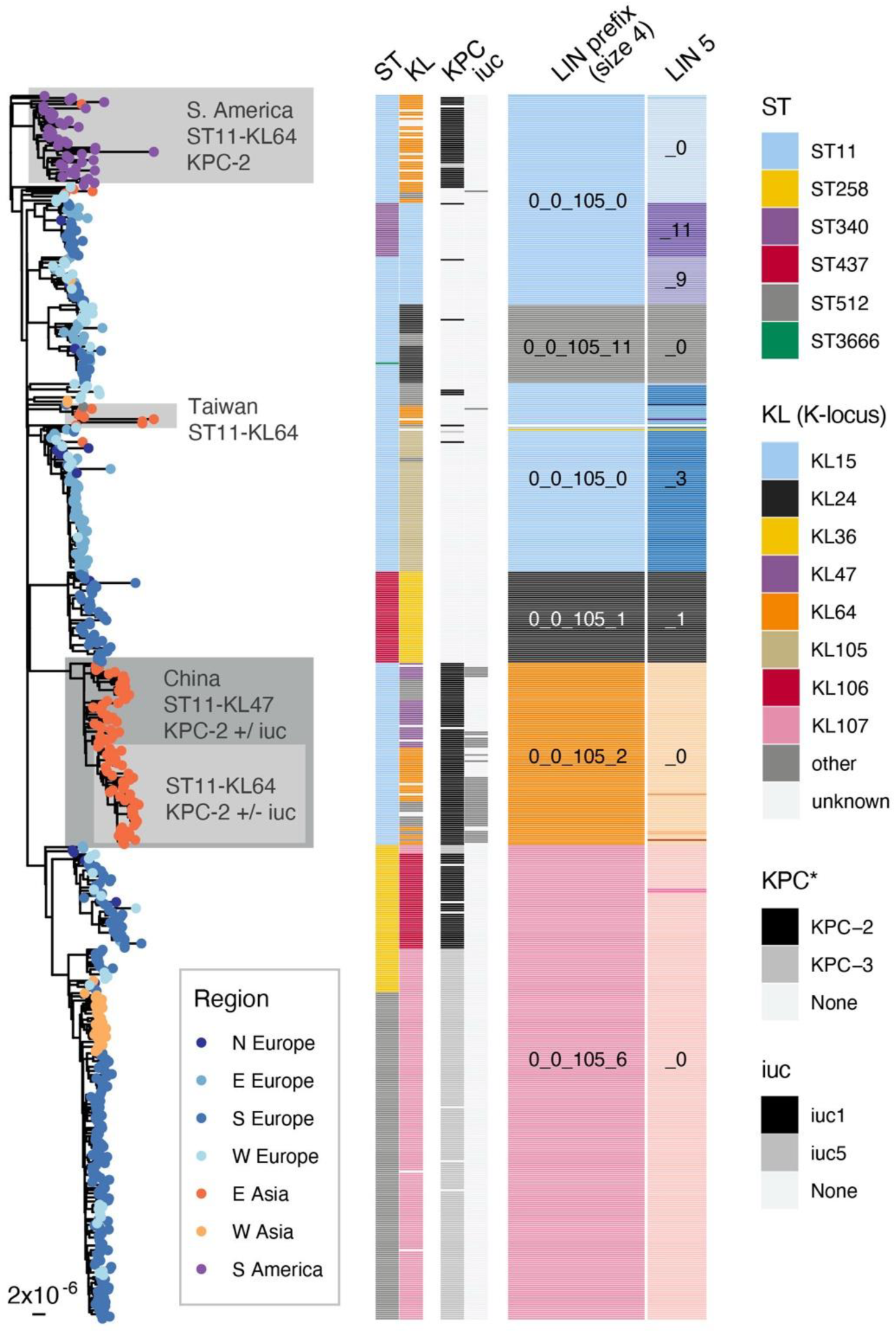
SL258 phylogenetic structure and LIN codes. Maximum likelihood phylogenetic tree of SL258 genomes inferred from a recombination-free variable site alignment (Supplementary appendix, **Section 12**). Tips are colored to indicate geographic regions of origin as per the legend (United Nations region classifications). The distribution of 7-gene multi-locus sequence types (STs), K-loci (KL), *bla*KPC (KPC variants), aerobactin locus lineages (*iuc*), and LIN code prefixes of sizes 4 and 5, are indicated by colored blocks as labelled (note that colors are independent to each column; for readability the labels for rare groups are omitted). Only K-loci identified with a Kaptive v2 confidence score of ‘Good’ or better are shown (otherwise marked ‘unknown’). Two isolates were detected with *bla*KPC-30 and one with *bla*KPC-12, but are not shown in the figure for brevity. Phylogenetic clades described in the text are colored and labeled accordingly.

LIN codes can help distinguish between different subclades that are associated with the same ST and capsule locus, a combination often used to describe specific subclones. For example, LIN codes clearly distinguish three phylogenetically distinct subclades that are all ST11-KL64 (grey shading on the tree branches, **Figure 5**). One of these is the major lineage circulating in China (0_0_105_**2**_**0_0_2**, predominantly 0_0_105_2_0_0_2_17) that carries KPC-2 and often the *iuc1* aerobactin virulence locus, as discussed broadly (36)(37). A second, unrelated ST11-KL64 subclade (0_0_105_**0**_**0**) is circulating in South America encoding KPC-2, but rarely *iuc* (38), while a third smaller clade (0_0_105_**0**_**2**) is detected primarily in Taiwan (39) rather than in mainland China (lacking KPC and with only one of eight genomes carrying *iuc*). These distinct clades are all referred to in the literature as ST11-KL64, despite representing phylogenetically distinct and likely unrelated, independently evolved, lineages. This example shows how LIN code classification beneath the sublineage level can help recognize and name subgroups of medical and epidemiological relevance, which should be subject to enhanced surveillance.

We next illustrate how LIN codes can subdivide isolates from single long-term outbreaks. Identifying outbreak strains and tracking strain diversification during outbreaks are key objectives of genomic epidemiology, as they provide capacity to quickly respond to outbreaks and prevent further infections. We use an Italian outbreak of *K. pneumoniae* SL147, a prominent multidrug-resistant international sublineage of *K. pneumoniae,* to show that clades that diversified during the outbreak are captured and labeled unambiguously with LIN codes (Supplementary appendix, **Section 13**). LIN codes were also recently used to label isolates from eight regional *K. pneumoniae* outbreaks that occurred within Poland (including two caused by SL258 and SL147), which had initially been loosely defined based on SNPs, O and K serotypes, and *bla*VIM carbapenemase-carrying integrons (40).

## Future directions and conclusions

Facilitating communication on the intra-species diversity of microbial strains is a key objective of strain taxonomies, which entail classification and naming of groups within species. In the field of epidemiological surveillance of bacterial pathogens, it has long been recognized that strain typing methods used for long-term and global strain tracking should be reproducible enough to enable internationally standardized nomenclatures, or ‘library typing systems’ (41). LIN codes based on cgMLST benefit from the high standardization and reproducibility of the cgMLST approach and provide a flexible and robust way to classify, name and identify subpopulations within bacterial species. The recognition of sub-populations associated with distinct phenotypes is an important “raison d’être” of taxonomies, and multilevel LIN codes strain taxonomies will advance our understanding of the links between genotypes and clinical phenotypes, vaccine coverage and antimicrobial resistance.

Given the reproducibility of cgMLST, an outbreak strain could be detected by different investigators based solely on its LIN code prefix. As LIN code prefixes are sufficient to define strain identity across countries or sectors, LIN codes provide a simple yet accurate solution for cross-border or other collaborative genomic surveillance investigations, without the need to share genomic sequences themselves (**Figure 6**). This possibility brought by shared strain taxonomies can alleviate issues around data confidentiality and sharing agreements, which are often an important barrier in genomic surveillance and rapid response to outbreaks under investigation by multiple institutional actors. Likewise, for the surveillance of particularly concerning strains, early warnings could be triggered based on the detection of the specific LIN codes of the targeted strains. LIN codes may thus become an integral part of epidemiological surveillance practice.

**Figure 6.**
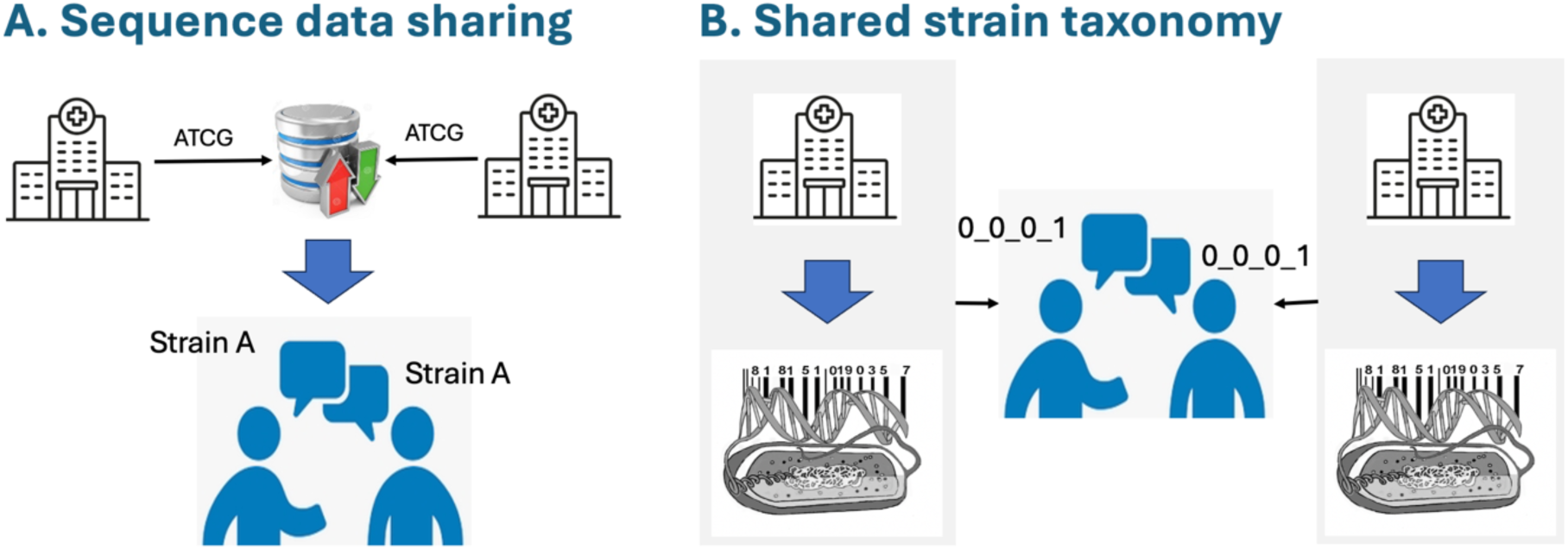
Two models of multicentric genomic epidemiology. **A**. The current model of sequence data sharing, which necessitates the sharing of sequence data from distinct institutions for their central analysis and comparison. **B**. The shared strain taxonomy model, where the use of a common nomenclature enables direct communication on subtypes and the recognition of identical strains without having to share sequence data.

LIN codes represent a widely applicable strain taxonomy system, as illustrated by the rapid pace of developments of LIN code implementations to bacterial pathogens. Following the initial use case for the KpSC, cgMLST LIN codes taxonomies have been introduced for *Streptococcus pneumoniae* (42), *Staphylococcus aureus* (43); *Moraxella catarrhalis* (44), *Neisseria gonorrhoeae* (45) and *Corynebacterium diphtheriae* (46). These LIN code taxonomies use different numbers of bins and thresholds adapted to the population structure of each species, illustrating the flexibility of the LIN code approach. The rapid development and adoption of LIN code taxonomies is facilitated by their integration into the BIGSdb platforms at Institut Pasteur and at Oxford University.

The applicability of the cgMLST LIN codes to most other bacterial species should be straightforward, provided they comprise sufficient genetic diversity. This requirement excludes the so-called monomorphic pathogens (47), such as *Mycobacterium tuberculosis* or *Salmonella enterica* serotype Typhi, where phylogeny-based taxonomies based on whole-genome SNPs are considered more useful given their higher resolution compared to cgMLST. The cgMLST LIN code strategy can also be extended with minor adaptations to other organisms with predominantly clonal reproduction, such as protozoan parasites and fungi, even if they are not haploid, given the existence of MLST taxonomies for *e.g., Candida albicans* and *Trypanosoma cruzi* (48,49).

The wide adoption of cgMLST LIN code strain taxonomies has the potential to result in a universal approach for standardized bacterial genotyping that could greatly enhance microbial biodiversity studies, international genomic epidemiology and infectious disease surveillance.

## Acknowledgements

We thank François Lebreton for providing the Newick file of the Italian ST147 outbreak described in Martin *et al*., 2021. We acknowledge the help of the HPC Core Facility of the Institut Pasteur for this work.

## Authors contributions

*Klebsiella* network genomic surveillance platform (KlebNET-GSP) conceptualization and coordination: SyB (Sylvain Brisse), KEH, DMA. cgMLST LIN code conceptualization and developments: MH, AC, SyB. BIGSdb-Pasteur platform maintenance: FP, BR, BB, SeB (Sebastien Bridel), SyB. Data acquisition and curation: VP, RI, CR, FP, MH, CC, SeB, ML, KLW, CAY, MRP, AC, DA. Data analyses: MH, SeB, KLW, ML, CAY. PubMLST and BIGSdb platform software development: KAJ, MCJM. Pathogenwatch platform maintenance and software development: DMA, CAY, SD. Data visualization: FP, MH, KLW, ML, SeB, CC, AC. Writing of first draft: SyB, with help from FP, MH, SC, KLW, KEH and AC. All authors contributed to, and approved, the final version of the manuscript.

## Ethical statements

Not relevant.

