## Supplementary appendix for "Life Identification Numbers: A bacterial strain nomenclature approach"

### Contents

#### Supplementary text (sections 1 to 13) and Figures S1 to S9


1. Formalization of the process of LIN code assignment
2. Impact of missing data
3. Comparison of LIN codes with the hierarchical clustering method
4. The importance of computational precision
5. Illustration of the impact of the input order on LIN codes
6. LIN code prefix trees tool: LINtree
7. LIN code taxonomy designed for the *Klebsiella pneumoniae* species complex
8. Nicknames attributed to LIN codes prefixes are inherited from 7-gene MLST up to ST6500
9. LIN code search and lookup functions in BIGSdb isolates databases
10. Use of LIN codes in external databases: the example of Pathogenwatch
11. Many Klebsiella pneumoniae 7-gene MLST sequence types are polyphyletic
12. Phylogenetic analysis of SL258: dataset and methods
13. Application of LIN codes to outbreak strain tracking: example of *K. pneumoniae* SL147


#### Supplementary references

#### 1. Formalization of the process of LIN code assignment

Formally, LIN codes are attributed to core genome Sequence Types (cgST) (1). Therefore, before assigning LIN codes, cgMLST profiles must be assigned to cgSTs. Like the ST designation in classical 7-gene MLST, a cgST is defined for each unique cgMLST profile, characterized by a unique combination of alleles at all loci of the scheme. Profiles with too many missing loci can be filtered out at this stage. A pair of cgSTs that differ only by missing data are defined as distinct cgSTs but correspond to a dissimilarity of 0% when ignoring missing data in cgMLST profile comparisons; these sets of cgSTs, here called 'coincidental cgSTs', are assigned the same LIN code (see next section).

The formal process of assigning LIN codes to cgSTs, based on their corresponding cgMLST profile, is described below:

> The LIN code of the first allelic profile is attributed 0 in every bin. Next, each new allele profile *j* is encoded from its closest already encoded profile *i* (*i.e.*, that maximizes the allele similarity percentage $s_{ij}$). After determining the pivot bin *p*, such that $s_{ij} \in [s_p, s_{p+1}[$ (*i.e.,* right threshold exclusive), the encoding of the new profile *j* is performed in three steps:
>
> (i) the same prefix as code *i* is attributed up to the bin $p-1$ (inclusive);
>
> (ii) for the pivot bin *p*: the maximum value observed in this bin among the subset of codes sharing the same prefix is incremented by 1;
>
> (iii) 0 is attributed at each downstream bin from $p+1$ (inclusive).
>
> Of note, when $s_{ij}$ = 100%, the LIN code of the new profile *j* is given the complete LIN code of *i* (including at the last bin).

#### 2. Impact of missing data

This section describes the impacts of missing data, leading to coincidental cgSTs and equal matches to several predefined LIN codes.

##### *2.1. Coincidental cgSTs and unique LIN codes*

Whereas 7-gene MLST genotyping requires complete allelic profiles to define a sequence type (ST), cgMLST approaches should tolerate the presence of missing alleles, as some core genes may not be essential, and as genome assembly shortfalls occasionally result in the absence or incompleteness of some loci. Therefore, the definition of cgSTs needs to accommodate missing data.

Profiles may differ only by loci where there is one or more missing allele(s) in one of the profiles, while otherwise identical at all loci called in both profiles. Such profiles will be assigned to distinct cgSTs. We define as coincident cgSTs, every group of cgST profiles that differ only by their missing data pattern (**Figure S1**). This may arise, in particular, when near-identical isolates or different sequencing runs of the same isolate lead to variable missing allele calls but are otherwise identical in the called loci, therefore leading to the creation of two or more coincident cgSTs.

As the dissimilarity between profiles is computed based solely on loci called in both profiles, each pair (or group of size >2) of coincident cgST profiles has a 0% dissimilarity value, and therefore the same LIN code.

When a given genome profile matches two or more predefined coincident cgSTs, it is attributed (by definition) to all the coincident cgSTs.

An isolate's profile may match ($d$=0) more than one encoded cgSTs, even if those are not coincident, due to missing loci in the isolates' genome (*i.e.*, at least one uncalled locus, at which two non-coincidental cgSTs have a distinct allele). In this case, a unique LIN code is defined (and displayed) for the isolate. To choose between the different possibilities, the LIN code of the cgST with the fewest missing allele(s) is attributed. When two or more coincident cgSTs have the same number of missing allele(s), the cgST with the smallest LIN code partition identifiers (considered from left to right bin, *i.e.*, the lowest sort order) is selected.

The same priority rule is applied to encode every novel profile that is equidistant (with $d > 0$) to two (or more) previously LIN-encoded non-coincidental cgSTs, in order to choose the reference LIN code from which the novel LIN code is created.

Note that if two genomes yield identical cgMLST profiles, including in their missing data pattern, they correspond to the same cgST.

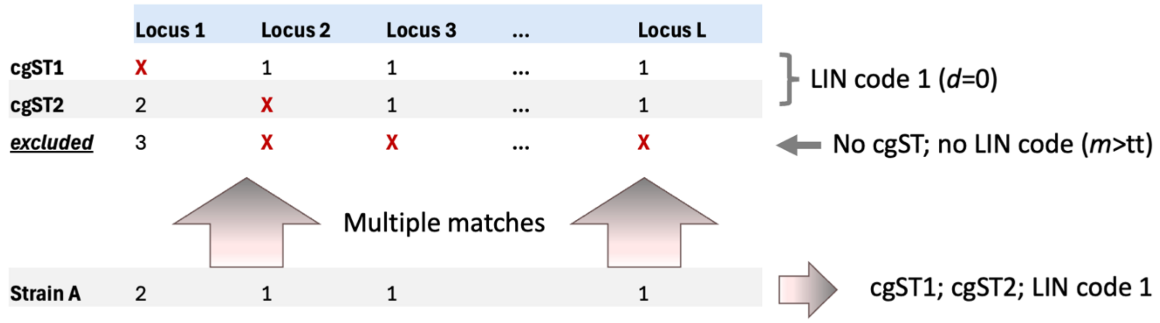


**Figure S1. Effect of missing data: coincidental cgSTs and multiple matches**

Core genome Sequence Types (cgST) are defined only if the number missing data $m$ is lower than the tolerance threshold ($tt$, set at 2 in this example). cgST1 and cgST2 are defined, as they have fewer missing data than $tt$. The third profile does not define a cgST, as $m > tt$. cgST1 and cgST2 are coincident cgSTs and then correspond to the same LIN code. Strain A matches both cgST1 and cgST2 and is attributed their unique LIN code.

*2.2. Definition of a maximal number of missing data*

Compatibility with missing data is an integral feature of cgMLST-based approaches (see above), but accepting too much missing data would lead to imprecise classifications. To minimize the complexities introduced by the above-described equal matches, a maximal number of accepted missing data (*e.g.*, 5% of the cgMLST scheme length) must be defined when implementing the cgST classification (**Figure S1**). Within BIGSdb, this value must be set when defining the LIN code scheme. cgMLST profiles can also be assigned to a cgST only when they comprise fewer than a defined (once for all) number of missing alleles (*i.e.*, profiles have a number of called alleles greater or equal to a predefined number).

For example, KpSC profiles with 30 (4.77%) or more missing alleles (likely to correspond to poor quality genomes) are not included in the KpSC LIN code taxonomy. For LIN code taxonomies in other bacterial species, the proportion of tolerated missing data for cgST assignment can be set to higher proportions (to increase the number of LIN-encoded genomes) or lower proportions (to improve the precision of LIN code classifications).

#### 3. Comparison of LIN codes with the hierarchical clustering method

Hierarchical clustering was developed by Mark Achtman's group and implemented in EnteroBase (2)(3). It was developed to address the well-known issue of instability of the classifications based on single-linkage clustering. Hierarchical clustering, implemented using the pHierCC pipeline, introduced a distinction between two phases in the process of genome encoding. First, a *development* mode, where a minimum spanning tree (MStree) approach (leading to endpoint results similar to those obtained with the single-linkage algorithm) is used to create groups. During this phase, MStrees are recalculated at each genome batch inclusion. Therefore, group fusion can occur, and the taxonomy is unstable and considered temporary. Second, a *production* mode, where individual genomes are successively added individually to the closest group without re-calculating the MStree. Static cluster assignment is thus achieved, without fusion of prior groups.

The genome encoding process in production mode is comparable to the LIN encoding process. However, whereas the switch to production mode is carried out only when the number of available genomes is judged sufficient to represent the population structure, the LIN codes assignment process is distinct in the way that LIN codes are attributed using a homogeneous process, starting from the first genome inclusion.

A second important difference lies in the fact that hierarchical clustering attributes partition numbers independently across levels, whereas LIN codes subdivisions of each upper partition (defined by an identical prefix) are attributed identifiers that start with 0. This difference leads to a predominance of small identifiers in LIN codes, whereas partition identifiers grow continuously and comprise large numbers in the lowest (rightmost) levels of hierarchical clustering. For example, there are currently >600,000 distinct group identifiers (integer values) at the shallowest HierCC levels for *Salmonella*, and over 300,000 for *E. coli* (https://enterobase.warwick.ac.uk, accessed July 28th, 2025), which implies the need to handle large numbers when communicating using HierCC taxonomies. The independence of group numbering at each level in HierCC also implies redundancy, as higher levels can be deduced by lower-level identifiers (but only by consulting the database schema as such relationships are not conveyed in the name). In contrast in the LIN code system, the initialization at 0 minimizes the redundancy in the resulting codes. An advantage of independent numbering as performed in hierarchical clustering is that partition numbers at any level are self-sufficient to describe groups, whereas LIN code partitions can only be understood in the context of their prefixes, which define their upper-level membership.

#### 4. The importance of computational precision

As for all categorizations that rely on thresholds, computational precision is critical for reproducible results. Here, the pairwise dissimilarity between cgMLST profiles, which is a ratio, may often have a higher number of decimals than can be handled by the computing system, and its rounded value may lead to a slight underestimate (or overestimate) of the true value. When the (true) dissimilarity between an incoming profile and its reference is identical to the left threshold of a bin (*i.e.*, the same ratio of distinct versus called alleles), a rounded value may incorrectly correspond to the previous bin (**Figure S2**). Therefore, pairwise dissimilarity computations should be performed with the same precision as the bin thresholds themselves. In BIGSdb, ratios corresponding to the thresholds are compared to the calculated dissimilarity values using Perl platform-native floating-point values (usually IEEE 754 double-precision).

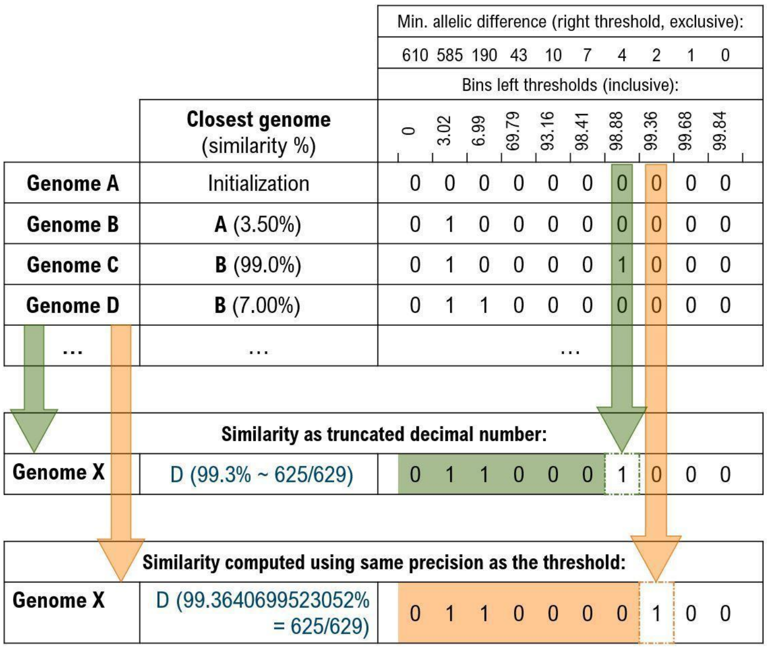


**Figure S2. The effect of rounded cgMLST similarity values on LIN code assignment.**

In this example, the use of a rounded value for the similarity between genome X and genome D leads to a slight underestimate, therefore creating a novel identifier in bin 7, instead of bin 8 when computing the similarity with the same precision as the threshold.

#### 5. Illustration of the impact of the input order on LIN codes.

The LIN code approach is dependent on input cgST order, as the partition in a given bin may vary slightly according to the order by which the genomes were encoded (1). We illustrate this effect on **Figure S3**.

To minimize this effect, BIGSdb uses the traversal of a minimum spanning tree (MStree) following Prim's algorithm (4) to define the order by which the novel profiles are encoded; this ensures that a minimal number of partitions are defined. This approach was implemented since BIGSdb v1.36.1 and also maximizes reproducibility when adding a batch of novel genomes. For each novel batch of genomes to be LIN-encoded, a MStree is created, and the isolate chosen as the starting point for LIN encoding is the one that has the closest similarity to an already encoded isolate in the database; next, the MStree is traversed from this node for encoding the remaining genomes. When creating LIN codes, novel genomes should ideally be encoded in batches as large as possible.

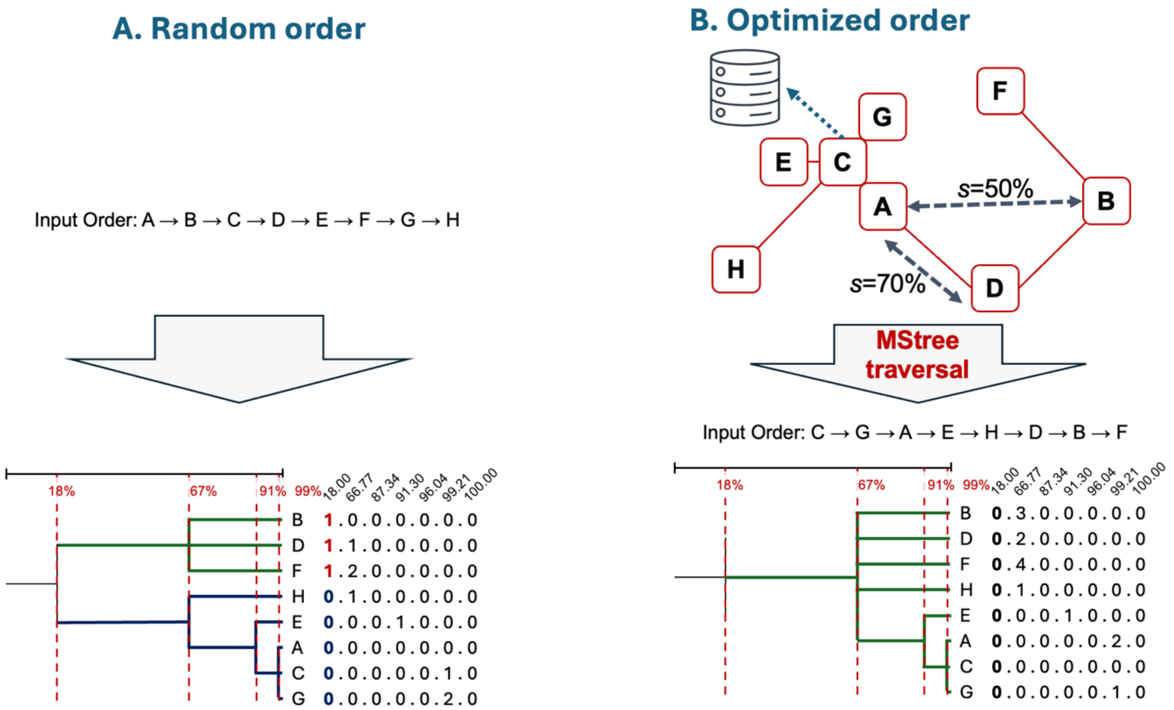


**Figure S3**. **Optimizing the input order for LIN code assignments.**

The resulting LIN code assignments can be affected by the input order of genomes into the taxonomy. **A**. Random order. **B**. With optimized order. When using a minimum spanning tree (MStree) to define the input order, the resulting number of partitions can be minimized, as following a MStree traversal order minimizes distance gaps between newly encoded genomes and those already present in the LIN code database. In this example, the MStree traversal avoids fragmentation of level 1, with a single partition (right) instead of two partitions (left) created by the early introduction of B. The difference is explained by the similarity of genomes A and B against genome D, which is higher than the bin 2 threshold (66.77%). In BIGSdb, the first isolate is also chosen to be the one with the shortest distance to the LIN code database members (genome C in this example).

#### 6. LIN code prefix trees tool: LINtree

LINtree (https://gitlab.pasteur.fr/GIPhy/LINtree) was developed and made available to build prefix trees. The input file contains a list of genome names and LIN codes (one sample per row), with a header row indicating the level of similarity for each bin. LINtree outputs a Newick-formatted tree showing the relationships between input genomes, based on the hierarchy provided by the LIN codes and with branch lengths scaled using the similarity levels in the header row. For example, the tree of the ST147 Italian outbreak (see corresponding section below) was generated using this tool, based on the input list of LIN codes. This example illustrates how the prefix tree recapitulates the phylogenetic relationships of this outbreak strain with its ancestral relatives, providing a useful aid in outbreak investigations.

#### 7. LIN code taxonomy designed for the *Klebsiella pneumoniae* species complex

KpSC profiles are defined using a 629-loci cgMLST scheme defined in the BIGSdb-Pasteur KpSC database: https://bigsdb.pasteur.fr/cgi-bin/bigsdb/bigsdb.pl?db=pubmlst_klebsiella_seqdef&page=schemeInfo&scheme_id=18. LIN codes are defined based on the variation recorded with this cgMLST scheme (database name: scgMLST629_S; common name: KpSC scheme), with 10 bins being defined. Bins 1 to 4 have as right thresholds 610, 585, 190 and 43 allele mismatches, respectively, while bins 5 to 10 correspond to thresholds 10, 7, 4, 2, 1 and 0 mismatches, respectively. Thus, the first bin corresponds to the range [629-610[ of cgMLST mismatches, whereas the last one corresponds to the range [1-0[. Note that the last bin excludes complete identity (*i.e.,* 0 mismatch*,* 629 matches); in this case, the LIN code is simply copied from the matching cgST.

To be encoded into the LIN code taxonomy, profiles need to have no more than 30 missing alleles out of the 629 loci.

The KpSC LIN code taxonomy is formally defined in BIGSdb-Pasteur as:

scheme id: 18 [scgMLST629_S]

levels: 610;585;190;43;10;7;4;2;1;0

max. missing alleles: 30

A total of ten bins were thus determined. The first four bins, which represent the deepest hierarchical levels of relatedness, correspond to species, subspecies, sublineage and clonal group, respectively (**Figure S4**). The last bins delineate six levels of high-resolution relatedness that might be useful for epidemiological surveillance. In this particular case, the cgMLST scheme is restricted to 629 loci, whereas a typical KpSC genome has over 5,500 genes, implying that the resolution is far from being complete, making the scheme more suited to public health surveillance. However, it can also be used to define clades during protracted outbreaks and new emerging strains (see examples of SL258 and SL147 in this manuscript).

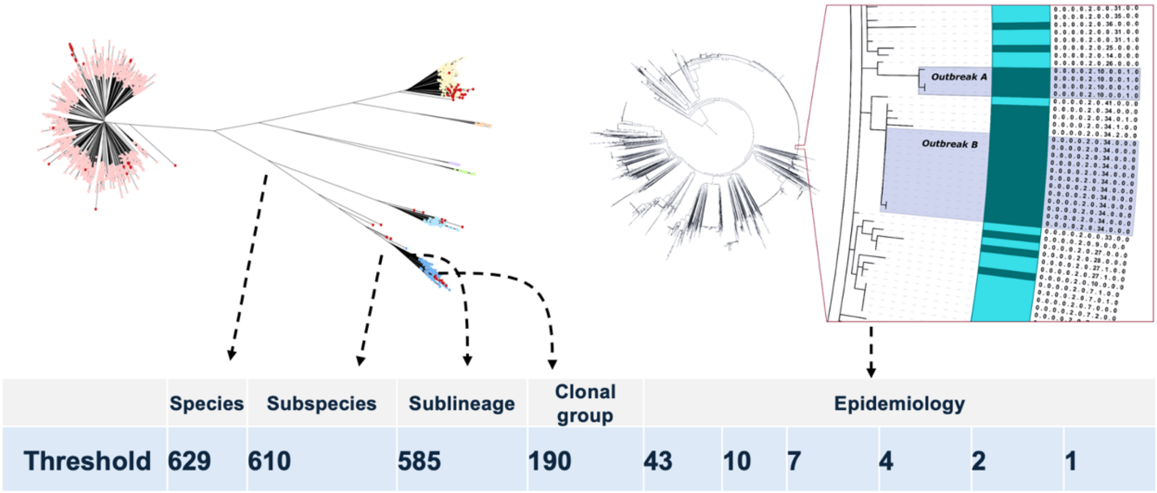


**Figure S4. The LIN code bin structure for the KpSC.**

The *Klebsiella pneumoniae* species complex (KpSC) LIN code system is based on a 629-loci cgMLST scheme. The 10 thresholds are given in the table as the maximal number of allele differences (inclusive) accepted in the corresponding bin. The four first (leftmost) bins were defined based on the population structure of the KpSC, whereas the 6 last bins, defined as a suite of decreasing thresholds (down to 1) are intended for use in epidemiological investigations.

#### 8. Nicknames attributed to LIN codes prefixes are inherited from 7-gene MLST up to ST6500

The MLST nickname inheritance rule was applied on LIN code prefixes of size 3 (sublineages) and 4 (clonal groups) as described in the main text. Note that the inheritance rule was applied only using ST identifiers up to ST6500. This captured the well-known major causes of KpSC human infections that have disseminated globally to-date. In order to make clear that the new nicknames are not inherited from MLST nomenclature for subsequent prefixes, SL and CG nicknames are numbered incrementally, starting with 10,000 (**Figure S5**).

In parallel, expansion of the MLST nomenclature will continue, resulting in defining novel STs (incremented by one) upwards of 6500 (currently the highest ST is ST9246, BIGSdb-Pasteur KpSC database, accessed October 26$^{th}$, 2025; comprising 9,221 profiles, as 25 STs were deleted and retired over the years, due to detection of errors *e.g.* in Sanger sequencing data). The correspondence that exists between ST identifiers > 6500 and prefix nicknames > 10,000 will not be immediately obvious but is available from the BIGSdb-Pasteur KpSC database.

Previously (1), cgMLST groups have been defined by the single-linkage (*slink*) clustering method using the same 10 thresholds as for the LIN codes, and the four highest-level groups were nicknamed by inheritance from Linnaean taxon names (for the two first) or MLST labels (for the levels defined by thresholds 190 and 43, dubbed Sublineage and Clonal Groups, respectively). Together with the LIN code taxonomy, which did not include nicknames at the time of the publication by Hennart *et al.* (1), this *slink*-based system formed a 'dual-barcoding approach'. However, because such *slink* groups suffered from fusion of existing groups upon addition of subsequent genotypes, which occasionally had intermediate distances between preexisting groups (*e.g.*, hybrid genotypes), the classification of cgMLST profiles into *slink* groups was abandoned. Fortunately, when excluding the hybrid genotypes, a nearly complete concordance was observed (1) at the four first levels between *slink* clusters and LIN code groupings (optimized by MStree traversal; see Input order section above). As a result, the LIN code taxonomy currently in use is nearly fully consistent with the one initially proposed in Hennart *et al.* (1), which has been abandoned. Only SL10000 to SL10021, and CG10000 to CG10276 correspond to groups that were renamed (table of correspondence available upon request). Rather than using a 'dual-barcoding approach' as proposed initially (1), the use of a single-barcoding taxonomic system, based solely on LIN codes, will stabilize and simplify the way KpSC groups are defined and labeled.

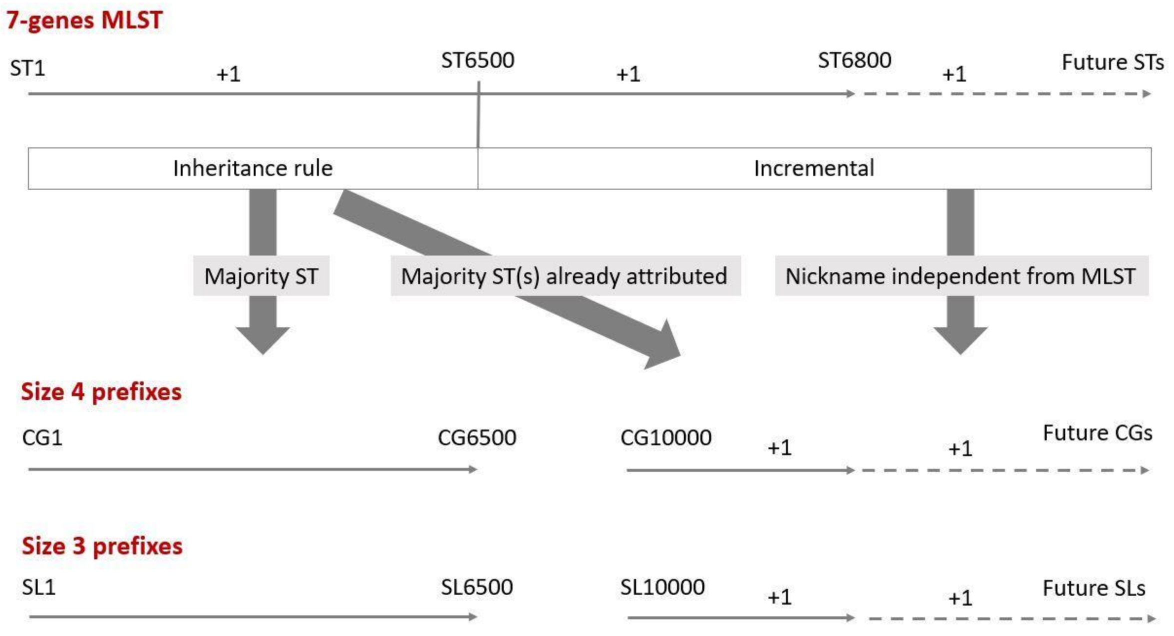


**Figure S5. LIN code prefix nicknames: application domain of inheritance from MLST.**

The top line represents the incremental definition of 7-genes MLST sequence types (ST). These identifiers up to ST6500 have been mapped onto LIN code prefixes of size 3 and 4, whereas for ST identifiers above 6500, no mapping was made and the prefix nicknames are incremented by 1, starting from 10,000. Future novel STs continue to be defined in parallel with no link to LIN code nicknames.

#### 9. LIN code search and lookup functions in BIGSdb isolates databases

Users can search for isolates of interest using the LIN code matching functionalities implemented within BIGSdb. A complete LIN code (or any prefix) can be used as a query (**Figure S6**). The nickname nomenclature attached to LIN code prefixes can also be used to facilitate the query of groups of interest (*e.g.,* SL258 members can be searched by using its attached prefix 0_0_105 or by using the SL258 nickname itself). Furthermore, related isolates at any level to a genome of interest can be looked up using the available LIN code function.

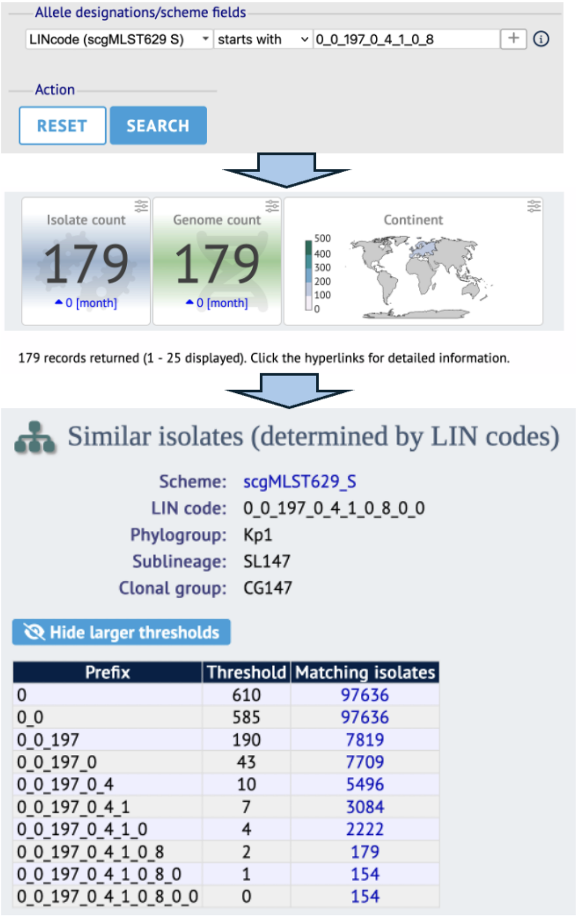


**Figure S6. Workflow to search for isolates with a given LIN code prefix.**

By clicking the Analysis menu item ‘Search by LIN code’, users may enter any prefix, choose ‘starts with’, enter a LIN code prefix, and click Search. Results will display all accessible isolates (public and private ones of the specific user). Isolate hyperlinks can then be clicked to display an isolate report page, where a lookup function for all isolates sharing any LIN code prefix is available.

#### 10. Use of LIN codes in external databases: the example of Pathogenwatch

As a pilot model of the external use of LIN codes, the KpSC LIN code taxonomy was implemented in the Pathogenwatch platform, which supports KpSC genomic typing (5) and in which allele, cgST and LIN code definitions are synchronized using the API functionality from BIGSdb-Pasteur into an internal temporary database. The cgMLST allele sequences extracted from the Pathogenwatch query genomes are compared to those in the temporary database, and the resulting query cgMLST profile is used to find the closest match in the temporary database. If the query genome does not match completely with an existing source nomenclature cgST, a provisional cgST is assigned, represented by an asterisk (*)

character followed by a temporary identifier (*e.g.,* cgST *f26e). Pathogenwatch also indicates the closest cgST defined in the source taxonomy database. Finally, a partially incomplete LIN code (containing asterisks in undefined bins) will be provided by Pathogenwatch based on the shared prefix with the closest reference cgST (**Figure S7**). This process provides information about the relatedness of query genomes compared to existing LIN code definitions. Given the good representativity of KpSC currently available genomic data for clinical isolates, this process can provide sublineage and clonal group identification in most cases. When Pathogenwatch provides provisional alleles, STs, cgSTs and/or LIN codes, the user should be encouraged to submit the genomic sequence data to the source BIGSdb-Pasteur database so that novel reference taxonomic elements (alleles, STs, cgSTs, LIN codes) can be created.

Inference of a query genome's LIN code in external resources can be carried out up to the bin preceding the pivot bin corresponding to the similarity with the closest reference match. In the KpSC system, when the LIN code prefix up to the fourth bin (*i.e.*, more than 586 shared alleles) can be defined for a query genome, information on species, SL and CG can be derived.

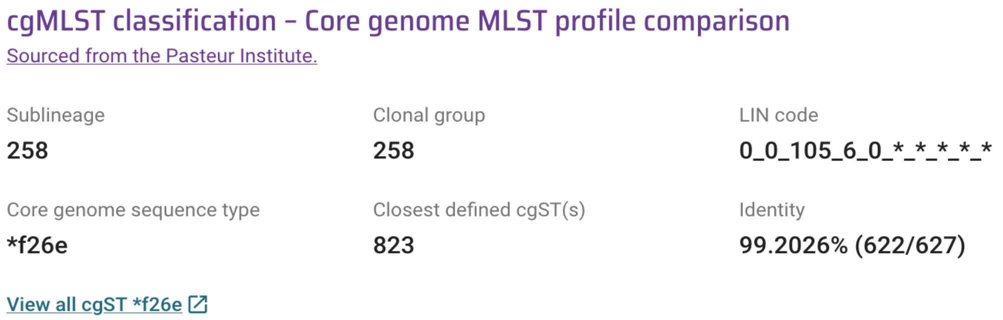


**Figure S7. Example of LIN code identification in Pathogenwatch.**
Although the LIN code is incomplete, the genome can be inferred to belong to sublineage SL258 (defined as prefix 0_0_105) and to clonal group 258 (prefix 0_0_105_6).

#### 11. Many *Klebsiella pneumoniae* 7-gene MLST sequence types are polyphyletic

Even though they are based on allelic profile comparisons rather than a sequence-based phylogenetic analysis, LIN code prefixes of length 3 or 4 bins are largely compatible with phylogenetic classifications and thus represent markers of their corresponding tree branches (1). In contrast, 7-gene MLST may conflate phylogenetically unrelated genomes in a single ST, for example through recombination leading to the same ST being assigned to genomes from distinct parental lineages, or by large recombination events affecting multiple cgMLST loci but leaving the 7-gene MLST loci unaffected (6).

Here we explore the extent of this phenomenon using 44,000 publicly available genomes of the *K. pneumoniae* species complex (accessed from BIGSdb-Pasteur KpSC database, June 2023). To spot potential discordances between ST and LIN code prefixes, we first filtered out non-Kp1 phylogroup (prefix 0_0) genomes and removed nearly identical cgMLST profiles, by keeping a single representative of each partition at LIN code level 5. STs observed only in a single isolate were then filtered out. We next searched for all STs that were split in several clonal groups or sublineages (as defined by their prefix) and conversely, also looked for prefixes of length 3 or 4 which comprised several STs. We then placed these genomes in a phylogenetic tree built using IQ-TREE v2.2.2.2 (7) using GTR+I+G model, from concatenated alignments of individual cgMLST gene alignments.

We found that 113 STs were polyphyletic, defined as being observed in at least two unrelated LIN code sublineages (**Table S1**). We illustrate this phenomenon for major STs in **Figure S8**. For example, ST485 was observed in four phylogenetically unrelated sublineages: SL485 (0_0_157), SL45 (0_0_158), SL1626 (0_0_227) and SL11569 (0_0_1215). This analysis also confirmed the polyphyletic status of ST23 (6), which conflates isolates from distant sublineages: SL23 (0_0_429) and SL218 (0_0_115) as illustrated on **Figure S8**.

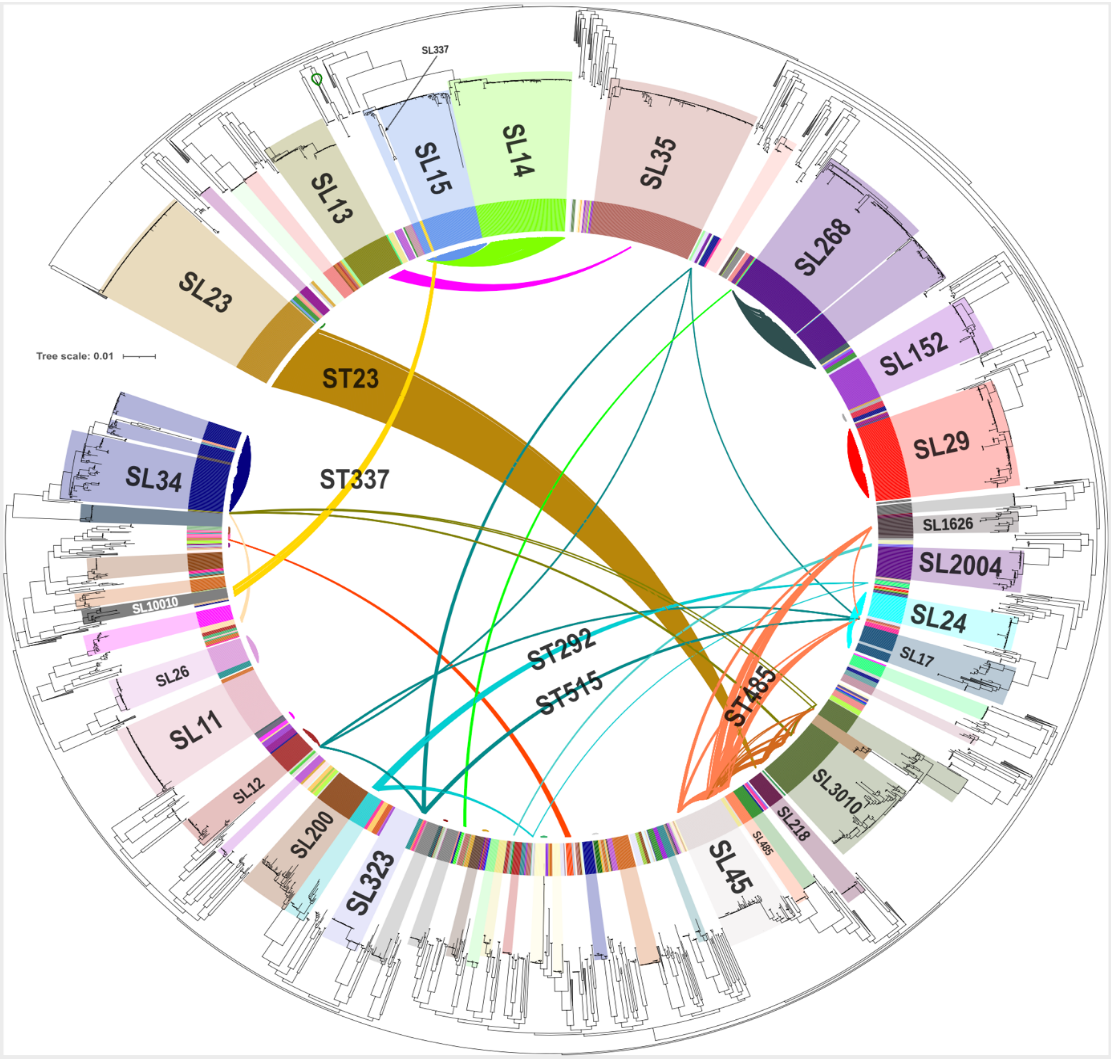


**Figure S8. Phylogenetic tree of *K. pneumoniae* main sublineages.**

A phylogenetic analysis of *K. pneumoniae sensu stricto* genomes (LIN code prefix 0_0; see **Table S2** for the list of included genomes) was performed from the multiple sequence alignments of 629 cgMLST genes. Closely related leaves were collapsed. The colored sectors in the inner circle correspond to the sublineages (SL) defined based on their LIN code prefix of length 3 (*i.e.*, made of the three first levels); the major sublineages are highlighted by lighter-colored sectors joining the circle to the tree leaves. The internal connectors between sublineages represent frequent STs that were found in two or more sublineages. The full interactive tree is available at: https://itol.embl.de/tree/1579917420525181688029926

#### 12. Phylogenetic analysis of SL258: dataset and methods

To illustrate the usage of LIN codes for defining genetic subdivisions within major sublineages of *K. pneumoniae*, the multidrug resistant sublineage SL258 was considered (see main text). For this purpose, whole genome sequences representing SL258 were identified among the EuSCAPE collection (8) and two recent studies reporting 7-gene ST11 with K-locus (KL) 47 and KL64, for which multiple independent evolutions have been reported (9,10). The ST11 genomes were subsampled to a manageable number as follows: (i) five randomly selected genomes per study year for each of ST11-KL47 and ST11-KL64 reported from sites across China, plus all ST11 with other K-loci reported in the same study included for context, totaling 92 genomes from this study (9); (ii) 64 genomes representing ST11-KL64 clade 1 as defined in an analysis of public ST11-KL64 genomes (10). Genome assemblies were acquired from Pathogenwatch and those with >500 contigs and/or total assembly size < 4,969,898 or > 6,132,846 bps were removed (as per the KlebNET-GSP quality control definitions). Kleborate v2.3.2 (11) was used to determine 7-gene ST, $\mathit{bla}_{\mathrm{KPC}}$ alleles, and *iuc* lineages (aerobactin locus), and Kaptive v2.0.7 (12,13) was used to identify KLs.

In order to infer a high-resolution phylogenetic tree, genome assemblies were used to simulate 100-bp paired end reads with wgsim (without errors, https://github.com/lh3/wgsim). Reads were mapped against the NJST258-1 completed reference genome (NCBI accession: CP006923.1) and single nucleotide variants called using the RedDog pipeline (https://github.com/katholt/RedDog). The resultant allele table was converted to a pseudo-whole genome alignment and used as input for Gubbins v2.3.2 (14), in order to detect and remove recombination (100 iterations). The final filtered alignment of 10,390 variable sites, representing 591 genomes, was used to infer a maximum likelihood (ML) phylogenetic tree using RAxML v8.2.9 with parameters: best of 5 runs, 1,000 bootstraps each, gamma model of rate variation (15). Subsequently, five genomes were removed due to excessive branch lengths. The ML tree was visualized with R v4.3.1 and the following packages: ape v5.7.1 (16), phytools v1.9-16 (17), and ggtree v3.8.2 (18).

#### 13. Application of LIN codes to outbreak strain tracking: example of *K. pneumoniae* SL147

SL147 is one of the most prominent multidrug-resistant international sublineages of *K. pneumoniae*, defined by its LIN code prefix 0_0_197.

**Figure S9** illustrates how the phylogenetic relationships within SL147 are captured by LIN codes, using a previously described dataset (19). SL147 comprises a single clonal group (0_0_197_0) and three 7-gene STs (ST147, ST273 and ST392). At LIN code position 5, four partitions (0_0_197_0_**0**, 0_0_197_0_**4**, 0_0_197_0_**17** and 0_0_197_0_**25**) correspond largely to ST273, ST392 and two deep branches of ST147. In addition, both ST147 and ST273 are genetically heterogeneous and structured phylogenetically into several minor branches, which were captured by additional partitions of LIN code level 5 (**Figure S9, panel A**).

Protracted outbreaks often lead their investigators to define local clades (or subgroups) within the closely related outbreak isolates. These clades are often attributed temporary placeholder names, which are difficult to compare across studies *e.g.*, Clade A and Clade B (20). We illustrate how LIN codes provide a way to define these clades definitively, using the diversity among outbreak isolates from the study of Martin *et al.* (20), corresponding to a protracted outbreak in Tuscany caused by a metallo-β-lactamase (NDM)–producing carbapenem-resistant ST147 (**Figure S9, panel B**). The dataset used is given in **Table S3.** The time span of the Tuscany outbreak is 2018 - 2021. Most of the isolates in this outbreak have prefix 0_0_197_0_**4_1_0**, thus differing by no more than 4 alleles (out of 629) with another member of the group. The authors defined two clades, A and B. Here, clade B corresponds to the set of LIN codes 0_0_197_0_**4_1_0_8_x_x** (*i.e.,* with prefix 0_0_197_0_**4_1_0_8**, with x meaning there may be variation at the two last positions). Clade A was more diverse, and LIN codes classify this genetic variability in a definitive way, with six $8^{th}$ position prefixes (0_0_197_0_4_1_0_**7**, 0_0_197_0_4_1_0_**9**, 0_0_197_0_4_1_0_**10**, 0_0_197_0_4_1_0_**11**, 0_0_197_0_4_1_0_**12** and 0_0_197_0_4_1_0_**66**). This example highlights how *K. pneumoniae* LIN codes can subdivide isolates from long-term outbreaks.

A search of the BIGSdb-Pasteur KpSC database (January $31^{st}$, 2024) for prefix 0_0_197_0_**4_1_0** identified n=395 *K. pneumoniae* genomes, isolated between 2014 and 2023 and coming from 20 countries from North America, Europe, Asia, Africa and Oceania, which indicate the global dissemination of this subgroup of SL147. However, prefix 0_0_197_0_**4_1_0_8** was so far only reported from the Italian outbreak. This example illustrates how LIN codes can facilitate the tracking of strain dissemination, by enabling the identification of similar isolates even though these were described in separate studies.

*Next page*:

**Figure S9. *Klebsiella pneumoniae* SL147 phylogenetic diversity and correspondence with LIN codes.**

**A.** Global diversity of ST147 (dataset from Rodrigues *et al*. 2022 Microb. Genomics 8 https://doi.org/10.1099/mgen.0.000737). The right part represents the complete LIN codes (bins 1 to 10), with blocks of colors highlighting identical prefixes.

**B**. Diversity of ST147 outbreak isolates in Italy (Martin *et al.* 2021 Proc. Natl. Acad. Sci. U. S. A. 118, e2110227118. https://doi.org/10.1073/pnas.2110227118). On the left, the phylogeny constructed based on whole-genome SNPs from Martin *et al.* is depicted (Newick tree file provided by the authors). On the right, is depicted the LIN prefix tree generated using LINtree (see below) based on the input list of LIN codes. Clade B defined in Martin *et al*. 2021 is characterized by a unique LIN code prefix (0_0_197_0_4_1_0_8).

437

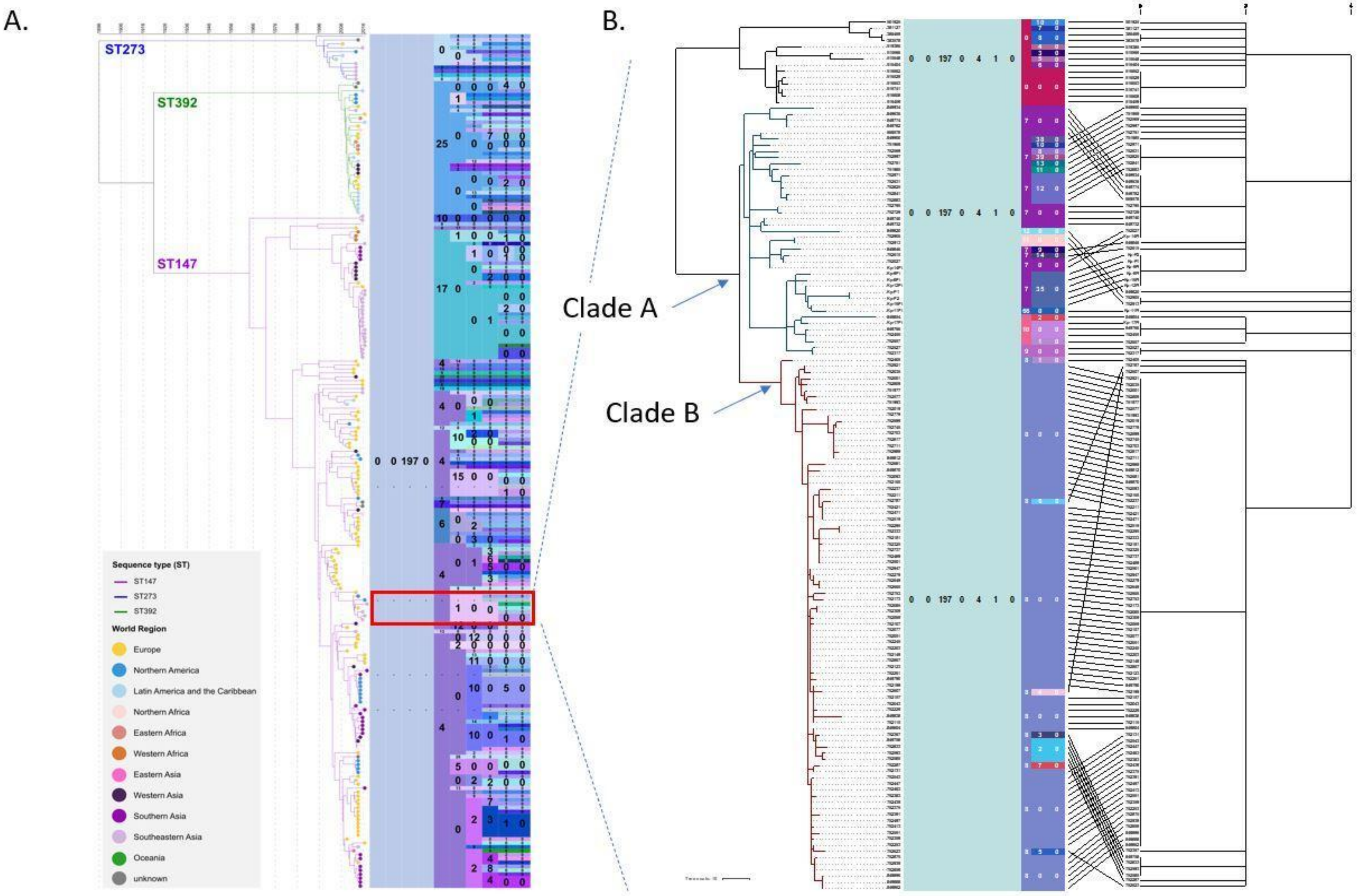
A.
ST273
ST392
ST147
Sequence type (ST)
ST147
ST273
ST392
World Region
Europe
Northern America
Latin America and the Caribbean
Northern Africa
Eastern Africa
Western Africa
Eastern Asia
Western Asia
Southern Asia
Southeastern Asia
Oceania
unknown
B.
Clade A
Clade B
